# Inverse PI by NMR: Analysis of Ligand ^1^H-Chemical Shifts in the Protein-Bound State

**DOI:** 10.1101/2021.11.30.470546

**Authors:** Gerald Platzer, Moriz Mayer, Jark Böttcher, Leonhard Geist, Julian E. Fuchs, Darryl B. McConnell, Robert Konrat

## Abstract

The study of protein-ligand interactions via protein-based NMR generally relies on the detection of chemical-shift changes induced by ligand binding. However, the chemical shift of the ligand when bound to the protein is rarely discussed, since it is not readily detectable. In this work we use protein deuteration in combination with [^1^H-^1^H]-NOESY NMR to extract ^1^H chemical shift values of the ligand in the bound state. The chemical shift perturbations (CSPs) experienced by the proton ligand resonances (free vs bound) are an extremely rich source of information on protein-ligand complexes. Besides allowing for the detection of intermolecular CH-π interactions and elucidation of the protein-bound ligand conformation, the CSP information can be used to analyse (de)solvation effects in a site-specific manner. In conjunction with crystal structure information, it is possible to distinguish protons whose desolvation penalty is compensated for upon protein-binding, from those that are not. Combined with the previously reported PI by NMR technique for the protein-based detection of intermolecular CH-π interactions, this method represents another important step towards the ultimate goal of Interaction-Based Drug Discovery.

## Introduction

The design of small molecules that bind target proteins with high specificity and affinity is based on the optimization of complementary interactions in the bound state versus the respective unbound solvated states. One crucial interaction in drug design is that between the π-electrons of aromatic ring-systems and aromatic or aliphatic CH containing groups.^1^ These so-called CH-π interactions are the most common type of non-covalent interaction in protein-ligand interactions.^2^ While they have relatively small individual interaction energies between -1.5 to -3 kcal,^3-7^ due to cooperative behavior,^8^ and minimal desolvation penalty,^9^ the cumulative effect of several CH-π interactions in protein-ligand binding events can result in a major contribution to binding energy.^10^ Often CH-π interactions are grouped under the rather broad term hydrophobic interactions.

The hydrogen donor of CH-π interactions in protein-ligand complexes can come from either the protein or ligand resulting in CH^protein^-π^ligand^ and CH^ligand^-π^protein^ interactions respectively. The hydrogen donors in proteins are either sp^2^-hybridized CH-groups of aromatic sidechains of Tyr, His, Phe and Trp,^11^ aliphatic sp^3^-hybridized (CH_3_, CH_2_) groups which exist in all natural amino acid sidechains or the protein backbone (α-CH).^12^ For aromatic sp^2^-hybridized CH-groups, the interaction is most favorable when the donor proton is positioned directly above the centroid of the aromatic acceptor ring-system (central CH-π H-bonds) at a distance (hydrogen to ring centroid) of ∼2.5 Å and has a calculated energy minimum of -2.46 kcal/mol as shown in theoretical works for the benzene dimer.^3-5^ For aliphatic, sp^3^-hybridized hydrocarbons like the benzene-methane pair the interaction energy is smaller (−1,45 kcal/mol) but the donor group ideal positioning remains directly above the aromatic ring-system center.^6, 7^ Unlike classical H-bonds, CH-π interactions are neither highly directional, nor strongly distance dependent,^13^ and hence still favorably contribute to binding when oriented above the periphery of a ring system rather than its center (peripheral CH-π H-bonds).^10^

We previously presented the PI by NMR methodology which can quantify the strength of CH-π interactions in protein-ligand complexes.^11^ The approach relies on the pronounced shielding effect exerted by aromatic ring-systems on interacting protons. Using a combination of amino acid labeling strategies and sensitive ^1^H-^13^C protein NMR spectroscopy we investigated the CH^protein^-π^ligand^ interactions of the protein sidechain CH-groups of Trp engaged in CH-π interactions with aromatic ligand moieties. We could show that the extent of proton chemical shift perturbations (CSPs) of interacting CH-donor groups can be directly correlated with energetically favorable interactions.

While the previously reported application of PI by NMR can be utilized for CH^protein^-π^ligand^ interactions through detection of protein hydrogen donors, the approach presented here allows for the detection of the inverse case, CH^ligand^-π^protein^ interactions, through the detection of ligand hydrogen donors. The labeling techniques used for the detection of interactions with aromatic ligand π-systems via protein aliphatic or aromatic sidechain hydrocarbons^14^ are not applicable for the inverse case as ^13^C isotope labeling of donor CH-groups present on small molecule ligands are technically challenging or too costly and therefore not a practical option for drug discovery programs. To overcome this, we have utilized protein deuteration and ligand based ^1^H-NMR spectroscopy to probe CH^ligand^-π^protein^ interactions between small molecule inhibitors of the N-methyltransferase NSD3 which bind to the methyl-lysine binding pocket of the PWWP1 domain.^15^

## Results and Discussion

In this work we present a fast and reliable detection method of bound ligand proton CSPs for the identification and optimization of favorable CH-π interactions between ligand hydrocarbons (CH^ligand^) and protein aromatic amino acids (π^protein^). The strategy combines replacement of non-exchangeable protons by deuteration, and ligand based ^1^H NMR spectroscopy to probe CH^ligand^-π^protein^ interactions in protein-ligand interaction sites. Protein deuteration has revolutionized NMR structural biology by greatly extending the molecular weight limit of protein systems amenable to NMR spectroscopy.^16^ Additionally, measurements in D_2_O solution eliminate exchangeable (amide and hydroxyl) protons thereby further simplifying the NMR spectra and reducing undesirable relaxation pathways. The combined effect of spectral simplification and improved relaxation properties make the NMR observation of ligands bound to protein targets feasible.^17-19^ In order to obtain atom-specific CSP data for each individual ligand proton, the assignments of ligand signals both in the free and protein-bound form are required. While this is technically straightforward for the unbound ligand (Figure 1 / red ^1^H-1D), assignment of the protein-bound state (Figure 1 / blue ^1^H-1D) is more challenging and usually involves [^13^C,^15^N]-filtered [^1^H,^1^H]-TOCSY/NOESY experiments.^20-22^ Here we utilize a combination of protein deuteration with a conventional [^1^H-^1^H]-NOESY experiment in order to correlate unbound ligand proton signals with the protein-bound state from a single experiment. This allows for a direct extraction of CSP data of ligand protons upon target binding. These chemical shift changes provide unique information about any formed CH^ligand^-π^protein^ interactions while additionally providing information about the conformation of the ligand in the bound state.

**Figure 1:**
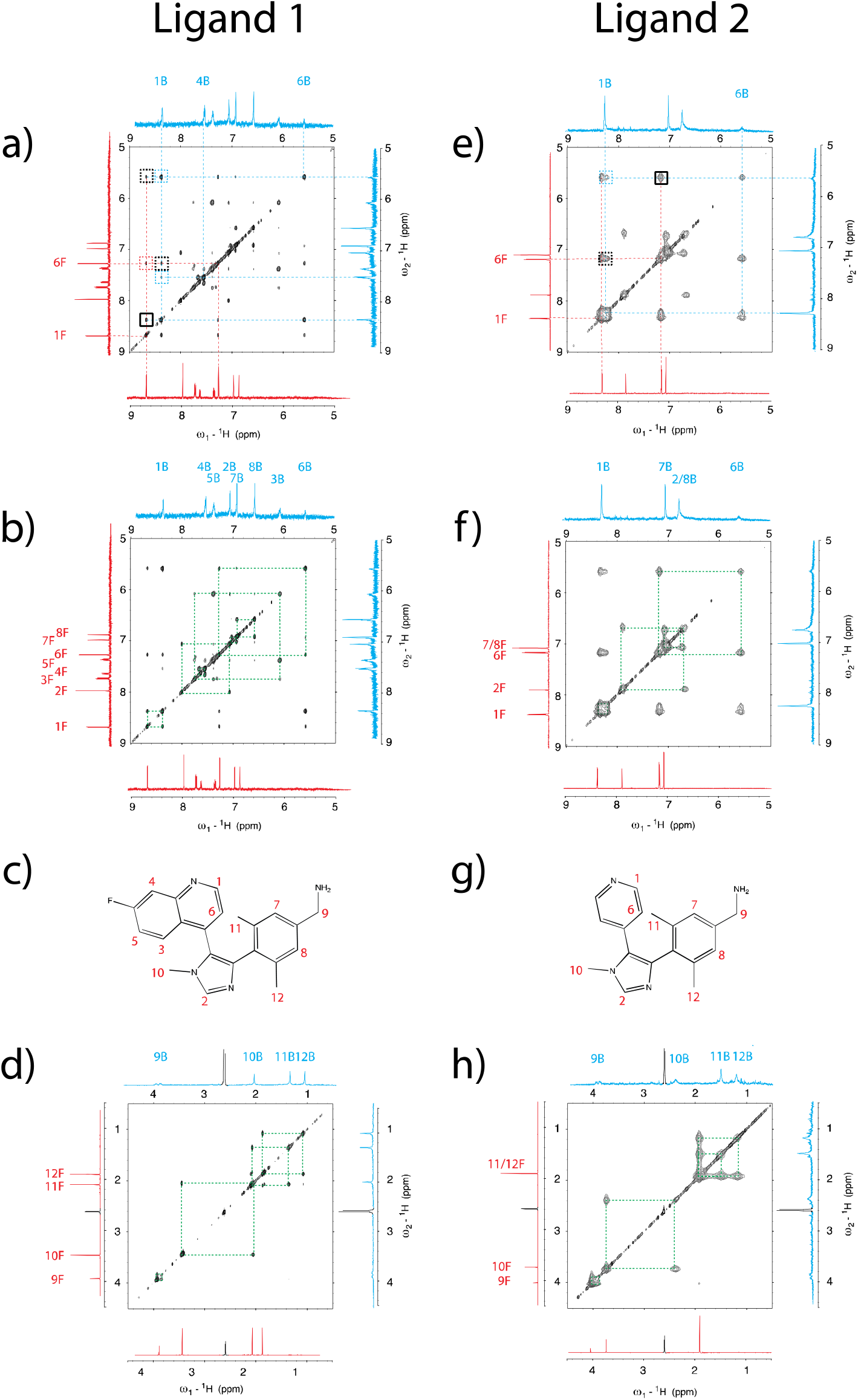
[^1^H-^1^H]-NOESY spectra of Ligand 1 (SPR KD = 170 nM) and Ligand 2 (SPR KD = 9.7 μM) in complex with the PWWP1 domain of NSD3. Spectra for the higher affinity Ligand 1 were recorded with a 2-fold excess over the protein, and with a 5-fold excess for the lower affinity Ligand 2. 1D traces represent ligand signals in the free state (red) and the bound state (blue). 1D peak suffixes F and B refer to the free and bound species respectively. [a), e)] Scheme for the assignment strategy of NOE cross peaks showing exchange peaks (solid black), exchange relayed NOE cross peaks (dotted black), NOEs between unbound signals (dotted red) and NOEs between bound signals (dotted blue). [b), f)] CSPs between free and bound signals (dotted green) for aromatic protons, and aliphatic protons [d), h)]. Individual structures of Ligands 1 and 2 are shown in [c), g)].

Originally the [^1^H-^1^H]-NOESY experiment was designed to probe spatial proximities of protons based on the r^-6^ dependence of the NOE effect. This information encoded in NOE cross peaks forms the basis for the extraction of distance constraints and is routinely used in NMR for structure determination. In drug design this is valuable information for medicinal chemists since it enables modeling of the ligand conformation when bound to the protein, especially in the absence of co-crystal structure information.^23^ Importantly for our Inverse PI by NMR strategy presented here, chemical exchange cross peaks between the free and bound forms can be also observed in the NOESY experiment (also known as EXSY) depending on the timescale of the binding kinetics. For binding processes in slow exchange, so-called exchange cross peaks arise from the reversible binding of the ligand to the protein during the mixing time of the NOESY experiment (usually in the order of 100 ms - 500 ms). This allows to link proton resonances of the free ligand with signals from the bound form, thereby providing direct access to ^1^H signal assignment of the protein-bound form. Assignment of the individual resonances is usually straightforward and can be carried out with three separate experiments using different ligand to protein ratios (see the SI for a discussion of the signal assignment strategy).

We chose the PWWP1 domain of the histone-lysine N-methyltransferase NSD3, a protein implicated in several forms of cancer, as the first system to demonstrate the methodology. The reasons for this choice were two-fold: the PWWP1 binding pocket of NSD3 is comprised of a cage of three potential aromatic π acceptor systems (Phe 312, Tyr 281 and Trp 284) and potent NSD3 inhibitors binding to the PWWP1 domain have been recently reported.^15^ The chemical probe molecule BI-9321 (named Ligand 1 here) (Figure 1c) binds to NSD3 with a SPR K_D_ of 170 nM and exhibits six positions (2, 3, 6, 10, 11 and 12) with both sp^3^- and sp^2^-hybridized CH donor groups which display significant upfield shifts (−0.70 to -1.74 ppm) between the unbound and protein-bound states in the [^1^H-^1^H]-NOESY experiments (Figure 2b). (Experimental vs. calculated ^1^H-CSP values for the individual positions of Ligand 1 are given in Table 1 of the SI) Large chemical shift perturbations are indicative of shielding effects of nearby aromatic ring-systems and indeed this is confirmed for five of the six (except position 2) upfield shifting CSPs upon inspection of the NSD3/Ligand 1 X-ray co-crystal structure (PDB ID: 6G2O) (Figure 2c). We have previously confirmed experimental vs calculated CSPs based on the crystal structure coordinates using the classical Pople formalism for CH^protein^-π^ligand^ interactions.^11^ In the case of the six NSD3 CH^ligand^-π^protein^ interaction CSPs observed, there is agreement between calculated and experimental results for CH donor positions 3, 11 and 12, while positions 2, 6 and 10 have an additional upfield chemical shift of around -1 ppm beyond what would be expected from aromatic ring-current anisotropy effects only. A similar behavior, although not explicitly discussed, is evident for the binding of GPI-1046 to the peptidyl-prolyl cis-trans isomerase FKBP12. ^24, 25^ For this system, the pyrrolidine protons embedded deep in the binding pocket experience a similar additional contribution to shielding that cannot be explained with the magnetic anisotropy arising from aromatic ring systems alone.

**Figure 2:**
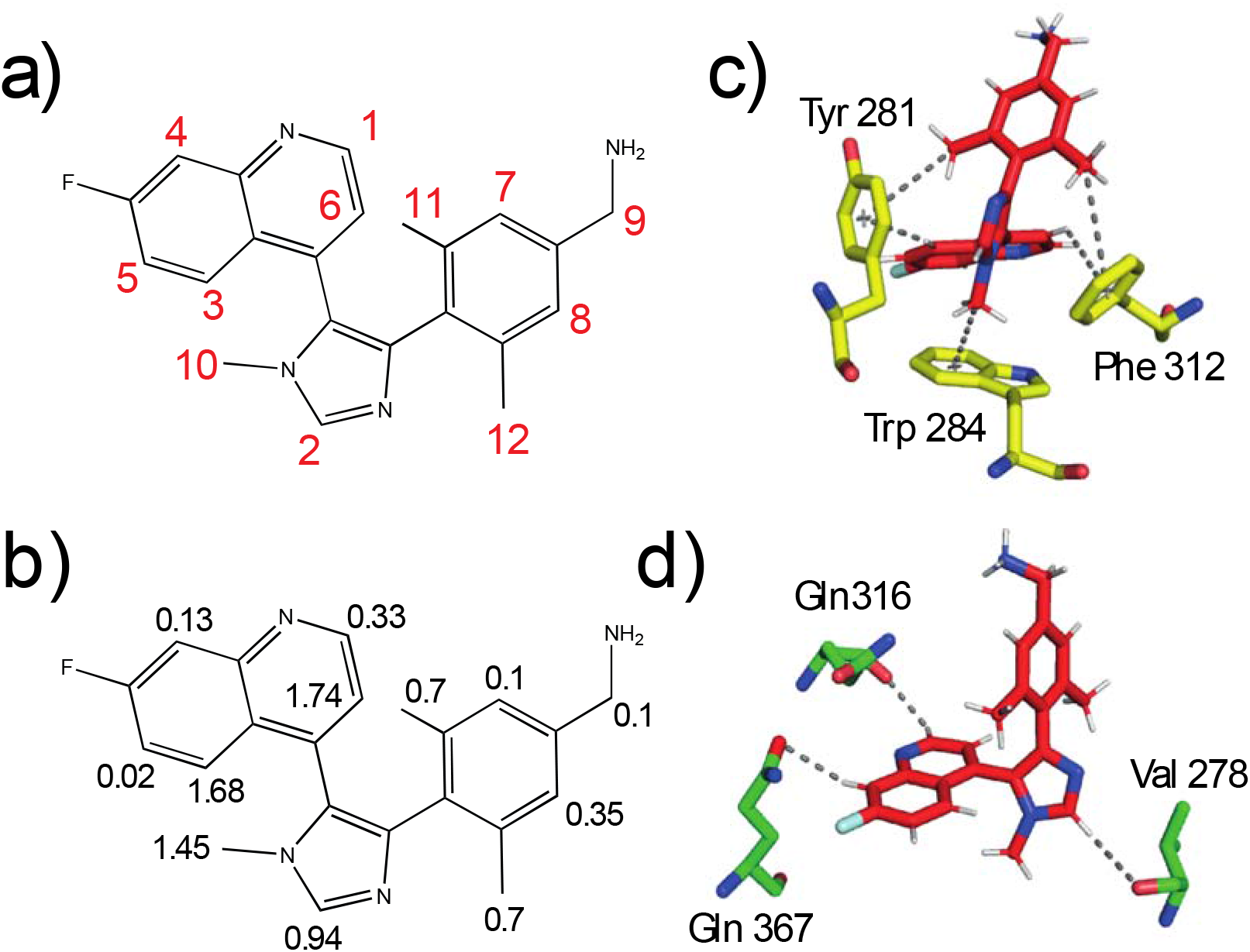
a) Proton signal numbering starting with the most deshielded signal. b) Observed chemical shift perturbations in ppm between the free and bound ligand. c) Contacts of bound Ligand 1 to aromatic residues in the binding pocket. d) CH-O contacts of Ligand 1 in complex with the NSD3-PWWP1 domain.

Positions 3 and 6 of the quinoline display the largest CSP upfield shift (around - 1.7 ppm) indicative of strong CH-π interactions. This is indeed confirmed by the crystal structure where position 3 forms a central CH-π interaction with the aromatic ring of Tyr 281 at a distance of 2.8 Å. For the quinoline proton at position 6 forms a peripheral CH-π interaction with Phe 312 which is 2.0 Å offset from the centroid. However, according to the Pople model, only -0.6 ppm upfield shift would be expected instead of the -1.74 ppm observed here.

The methyl group protons at positions 11 and 12 show upfield CSPs of -0.7 ppm upon binding to NSD3 which are in line with the Pople calculated values (−0.3 to -0.4 ppm). It is important to note that the CSP perturbation for methyl protons is approximately one third lower versus an individual CH interaction due to averaging of the three rotameric states of the methyl group protons. These interactions are confirmed by crystal structure information which shows that the methyl group of position 12 forms a peripheral CH-π interaction with Tyr 281 (distance 3.3 Å and offset 1.5 Å to centroid) while the position 11 methyl forms a peripheral interaction with Phe312 (distance 3.1 Å and offset 1.6 Å to centroid).

It is the imidazole CH donors for which the expected upfield CSP versus observed show the largest deviation. The imidazole N-methyl group (position 10) experiences an upfield CSP of -1.45 ppm upon NSD3 binding and a peripheral CH-π interaction with Trp 284 (distance 3.0 Å and offset 1.4 Å to centroid) is observed in the crystal structure. However, this configuration only results in a -0.46 ppm upfield CSP from the calculated ring current anisotropy using the Pople formalism. The most striking observation is the CSP of ligand proton 2. Here, no CSP would be expected for the imidazole CH (position 2) due to aromatic ring-current anisotropy as no CH-π interactions are evident in the 1.8 Å crystal structure of Ligand 1 bound to NSD3 and yet an upfield CSP of -0.94 ppm is observed.

The question arises what the origin of this additional shielding effect for some of these protein-bound CH groups is. The answer lies in the hydrogen-bonding properties of CH groups. Carbon-bound protons can, just like protons bound to more electronegative atoms like N or O, act as donor groups in hydrogen bonding interactions (e.g. CH-O, CH-π).^26-30^ In the context of ligand binding, CH-O interactions are of particular interest when considering hydrogen bonding interactions of CH, CH_2_ and CH_3_ groups with the solvent involving water as acceptor.^31^ Especially the participation of aromatic CH-groups in hydrogen bonding events with the solvent is often neglected.^32, 33^ Even though such hydrogen bonding interactions to the solvent are relatively weak and highly transient,^34^ the presence of a CH^ligand^-O^water^ hydrogen bond leads to a deshielding of the involved CH proton and hence a downfield shift in the NMR spectrum.^35^ This downfield shift can reach up to +1.5 ppm depending on the acidity of the donor protons.^36^ If in the protein bound state a proton is no longer solvated, this deshielding effect is relieved and an apparent upfield CSP is observed. This effect can partially be compensated by interaction with protein oxygens (O^protein^). In the case of Ligand 1, protons 1 and 4 are well compensated via the formation of CH^ligand^-O^protein^ H-bond with distances of 2.5 Å and 2.7 Å, respectively (Figure 2c). This compensation is well reflected in their small CSP values. Similarly small values (−0.02 to -0.35 ppm) are found for positions 5, 7, 8 and 9 all of which still have contacts to the solvent in the bound state and hence a minimal change to their experienced chemical shielding. For the imidazole proton 2, a CSP of -0.94 ppm suggests only a partial compensation which can be reasoned with the suboptimal hydrogen bond formed with the carbonyl oxygen of Val 278 and the high relative acidity of the imidazole CH group. Equally the protons in the positions 6 and 10, both with an additional upfield CSP of -1 ppm are not compensated via a CH^ligand^-O^protein^ interaction.

To widen the scope of this analysis to weaker affinity ligands often dealt with in the early stages of the drug discovery programs, we attempted this analysis with Ligand 2 which has an affinity of 9.7 µM (PDB ID: 6G2E) (Figure 1g).^15^ In contrast to the high-affinity system, only population averaged resonances are observed in the 2D [^1^H-^1^H]-NOESY spectra (Figure 1e). Importantly, even in cases of exchange line-broadening the sparsely populated (invisible) apo-state (unbound) is detectable in the NOESY spectrum. This is reminiscent of chemical exchange saturation transfer (CEST) type experiments.^37^ The EXSY pattern for the weaker binding Ligand 2 is almost congruent with the more decorated nM Ligand 1 for the chemical moieties shared between the two. This suggests that meaningful information about the presence of distinct CH-π interactions can be obtained in early stages of a drug development program even for weak-to-moderate binding affinities.

In addition to valuable CSP information, the NOESY experiment also gives access to distance information of the ligand-bound state encoded in NOE cross peaks, reporting on ligand protons in close proximity to each other. Figure 3a shows NOESY cross peaks (dotted blue) from ligand proton 3 to bound ligand signals 5, 10 and 12. Figure 3b shows the corresponding spatial proximity information directly mapped onto the crystal structure of NSD3 in complex with Ligand 1. The thereby extracted distance information provides insight into the relative positioning of ligand protons to each other, which can be especially relevant to determine the bound conformation of flexible ligands. Although this is not the prime motivation of our approach, the easy access to additional structural information about the bound conformation of the ligand can additionally support drug design efforts.

**Figure 3:**
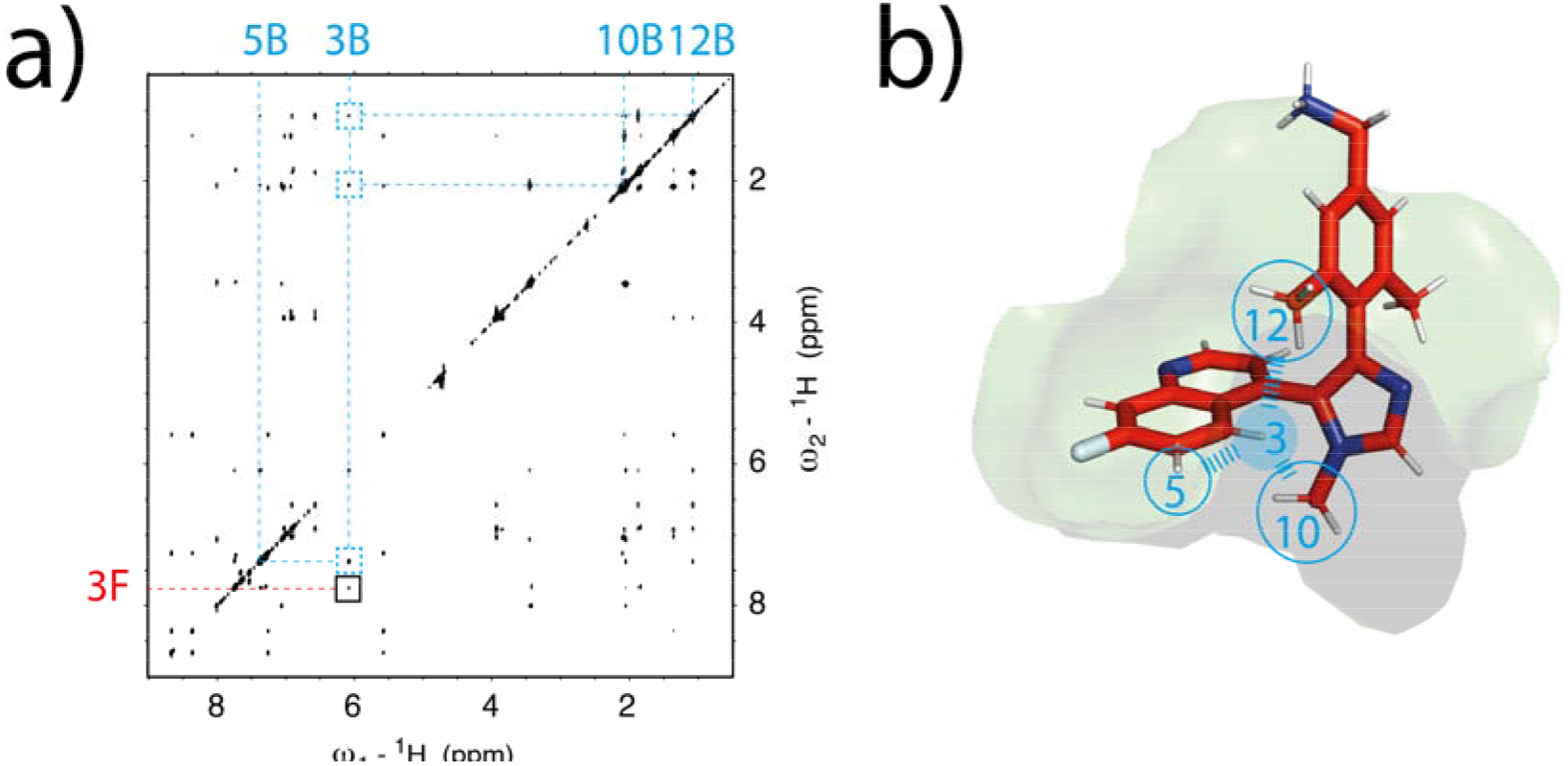
a) [^1^H-^1^H]-NOESY spectra of Ligand 1 (SPR KD = 170 nM) in 2-fold excess over protein. NOE cross peaks originating from proton 3 link to the spatially close proton signals 5, 10 and 12 (dotted blue). In addition, the exchange cross peak links to the free signal of proton 3 (solid black). The suffixes F and B refer to the free and bound species respectively. b) Bound state conformation of Ligand 1 complexed with NSD3-PWWP1 showing NOE extracted contacts for proton 3 to spatially close protons.

## Results and Discussion

We present a straightforward workflow to determine CSP data for ligand protons when bound to their target protein under saturating conditions. A wealth of information can be gained by careful analysis of the ligand’s free vs. bound CSPs, which have been largely untapped until now. While the analysis presented here is by no means exhaustive, our data suggests that we can contribute valuable information to important interactions such as CH-π, (de)solvation and its compensation in the bound state. We are not completely independent of a protein-ligand model or a complex structure at this point but are working on additional experiments to perform this type of analysis also in absence of a high-resolution crystal structure and on a timescale that is competitive in the fast-moving environment of medicinal chemistry. Thus, we have complemented the protein-derived PI by NMR approach described in our previous report,^11^ with a ligand-based version (inverse PI by NMR), that allows to visualize CH donors on the side of the ligand. This will allow a more comprehensive mapping of relevant CH-π interactions in the binding pocket. Large upfield ^1^H chemical shift changes in the ligand are induced by aromatic sidechains of the protein, provided that an energetically favorable CH-π interaction exists. In addition, our data suggests, that desolvation of CH groups that are not compensated by either forming a CH-π or a CH-O interaction in the bound state can also lead to significant loss of deshielding with CSP values of up to -1 ppm. In addition, we could show that even in the early stages of drug discovery programs, where ligands with moderate affinities need to be optimized, the (ligand-based) observation of chemical-shift changes induced by aromatic ligand moieties can be utilized to identify fragments which form the best CH-π interactions.

We anticipate that the implementation of both ligand- and protein-detected PI by NMR strategies will provide novel opportunities in future drug design, particularly for difficult-to-drug targets, by offering direct access to relevant molecular interactions in the protein ligand interface and strategic guidance for medicinal chemistry – both of which have the potential to change the way drugs will be designed in the future.

## Acknowledgements

G. Platzer was funded by the Christian Doppler Laboratory for High-Content Structural Biology and Biotechnology, Austria. The financial support by the Austrian Federal Ministry for Digital and Economic Affairs and the National Foundation for Research, Technology and Development is gratefully acknowledged.

## Notes

### Competing Interest Statement

The authors have declared no competing interest.

## References

1. Nishio, M.; Umezawa, Y.; Fantini, J.; Weiss, M. S.; Chakrabarti, P., CH-pi hydrogen bonds in biological macromolecules. Phys Chem Chem Phys 2014, 16 (25), 12648–83.

2. Ferreira de Freitas, R.; Schapira, M., A systematic analysis of atomic protein–ligand interactions in the PDB. MedChemComm 2017, 8 (10), 1970–1981.

3. Tsuzuki, S.; Honda, K.; Uchimaru, T.; Mikami, M.; Tanabe, K., Origin of attraction and directionality of the pi/pi interaction: model chemistry calculations of benzene dimer interaction. J Am Chem Soc 2002, 124 (1), 104–12.

4. Sinnokrot, M. O.; Valeev, E. F.; Sherrill, C. D., Estimates of the ab initio limit for pi-pi interactions: the benzene dimer. J Am Chem Soc 2002, 124 (36), 10887–93.

5. Podeszwa, R.; Bukowski, R.; Szalewicz, K., Potential energy surface for the benzene dimer and perturbational analysis of pi-pi interactions. J Phys Chem A 2006, 110 (34), 10345–54.

6. Ringer, A. L.; Figgs, M. S.; Sinnokrot, M. O.; Sherrill, C. D., Aliphatic C-H/pi interactions: Methane-benzene, methane-phenol, and methane-indole complexes. J Phys Chem A 2006, 110 (37), 10822–8.

7. Tsuzuki, S.; Honda, K.; Uchimaru, T.; Mikami, M.; Tanabe, K., The Magnitude of the CH/π Interaction between Benzene and Some Model Hydrocarbons. Journal of the American Chemical Society 2000, 122 (15), 3746–3753.

8. Ran, J.; Wong, M. W., Saturated hydrocarbon-benzene complexes: theoretical study of cooperative CH/pi interactions. J Phys Chem A 2006, 110 (31), 9702–9.

9. Scheiner, S.; Kar, T., Effect of solvent upon CH…O hydrogen bonds with implications for protein folding. J Phys Chem B 2005, 109 (8), 3681–9.

10. McConnell, D. B., Biotin’s Lessons in Drug Design. Journal of Medicinal Chemistry 2021, 64 (22), 16319–16327.

11. Platzer, G.; Mayer, M.; Beier, A.; Bruschweiler, S.; Fuchs, J. E.; Engelhardt, H.; Geist, L.; Bader, G.; Schorghuber, J.; Lichtenecker, R.; Wolkerstorfer, B.; Kessler, D.; McConnell, D. B.; Konrat, R., PI by NMR: Probing CH-pi Interactions in Protein-Ligand Complexes by NMR Spectroscopy. Angew Chem Int Ed Engl 2020, 59 (35), 14861–14868.

12. Plevin, M. J.; Bryce, D. L.; Boisbouvier, J., Direct detection of CH/π interactions in proteins. Nature Chemistry 2010, 2, 466.

13. Tsuzuki, S.; Fujii, A., Nature and physical origin of CH/pi interaction: significant difference from conventional hydrogen bonds. Phys Chem Chem Phys 2008, 10 (19), 2584–94.

14. Lichtenecker, R. J.; Schörghuber, J.; Bisaccia, M., Synthesis of Metabolic Amino acid Precursors: Tools for Selective Isotope Labeling in Cell-Based Protein Overexpression. Synlett 2015, 26 (19), 2611–2616.

15. Bottcher, J.; Dilworth, D.; Reiser, U.; Neumuller, R. A.; Schleicher, M.; Petronczki, M.; Zeeb, M.; Mischerikow, N.; Allali-Hassani, A.; Szewczyk, M. M.; Li, F.; Kennedy, S.; Vedadi, M.; Barsyte-Lovejoy, D.; Brown, P. J.; Huber, K. V. M.; Rogers, C. M.; Wells, C. I.; Fedorov, O.; Rumpel, K.; Zoephel, A.; Mayer, M.; Wunberg, T.; Bose, D.; Zahn, S.; Arnhof, H.; Berger, H.; Reiser, C.; Hormann, A.; Krammer, T.; Corcokovic, M.; Sharps, B.; Winkler, S.; Haring, D.; Cockcroft, X. L.; Fuchs, J. E.; Mullauer, B.; Weiss-Puxbaum, A.; Gerstberger, T.; Boehmelt, G.; Vakoc, C. R.; Arrowsmith, C. H.; Pearson, M.; McConnell, D. B., Fragment-based discovery of a chemical probe for the PWWP1 domain of NSD3. Nat Chem Biol 2019, 15 (8), 822–829.

16. Sattler, M.; Fesik, S. W., Use of deuterium labeling in NMR: Overcoming a sizeable problem. Structure 1996, 4 (11), 1245–1249.

17. Seeholzer, S. H.; Cohn, M.; Putkey, J. A.; Means, A. R.; Crespi, H. L., NMR studies of a complex of deuterated calmodulin with melittin. Proc Natl Acad Sci U S A 1986, 83 (11), 3634–8.

18. Fesik, S. W., NMR-Studies of Molecular-Complexes as a Tool in Drug Design. Journal of Medicinal Chemistry 1991, 34 (10), 2937–2945.

19. Hsu, V. L.; Armitage, I. M., Solution structure of cyclosporin A and a nonimmunosuppressive analog bound to fully deuterated cyclophilin. Biochemistry 1992, 31 (51), 12778–84.

20. Otting, G.; Wuthrich, K., Heteronuclear filters in two-dimensional [1H,1H]-NMR spectroscopy: combined use with isotope labelling for studies of macromolecular conformation and intermolecular interactions. Q Rev Biophys 1990, 23 (1), 39–96.

21. Breeze, A., Isotope-filtered NMR methods for the study of biomolecular structure and interactions. Progress in Nuclear Magnetic Resonance Spectroscopy - PROG NUCL MAGN RESON SPECTROS 2000, 36, 323–372.

22. Orts, J.; Gossert, A. D., Structure determination of protein-ligand complexes by NMR in solution. Methods 2018, 138-139, 3–25.

23. Erlanson, D. A.; Davis, B. J.; Jahnke, W., Fragment-Based Drug Discovery: Advancing Fragments in the Absence of Crystal Structures. Cell Chem Biol 2019, 26 (1), 9–15.

24. Sich, C.; Improta, S.; Cowley, D. J.; Guenet, C.; Merly, J. P.; Teufel, M.; Saudek, V., Solution structure of a neurotrophic ligand bound to FKBP12 and its effects on protein dynamics. Eur J Biochem 2000, 267 (17), 5342–55.

25. Wang, B.; Raha, K.; Merz, K. M., Jr., Pose scoring by NMR. J Am Chem Soc 2004, 126 (37), 11430–1.

26. Taylor, R.; Kennard, O., Crystallographic evidence for the existence of CH-O, CH-N and CH-Cl hydrogen bonds. Journal of the American Chemical Society 1982, 104 (19), 5063–5070.

27. Allerhand, A.; Von Rague Schleyer, P., A Survey of C-H Groups as Proton Donors in Hydrogen Bonding. Journal of the American Chemical Society 1963, 85 (12), 1715–1723.

28. Auffinger, P.; Louise-May, S.; Westhof, E., Hydration of C-H groups in tRNA. Faraday Discuss 1996, (103), 151–73.

29. Desiraju, G. R., Hydrogen bridges in crystal engineering: interactions without borders. Acc Chem Res 2002, 35 (7), 565–73.

30. Yesselman, J. D.; Horowitz, S.; Brooks, C. L., 3rd; Trievel, R. C., Frequent side chain methyl carbon-oxygen hydrogen bonding in proteins revealed by computational and stereochemical analysis of neutron structures. Proteins 2015, 83 (3), 403–410.

31. Panigrahi, S. K.; Desiraju, G. R., Strong and weak hydrogen bonds in the protein-ligand interface. Proteins 2007, 67 (1), 128–141.

32. Li, S.; Cooper, V. R.; Thonhauser, T.; Puzder, A.; Langreth, D. C., A density functional theory study of the benzene-water complex. J Phys Chem A 2008, 112 (38), 9031–6.

33. Slipchenko, L. V.; Gordon, M. S., Water-benzene interactions: an effective fragment potential and correlated quantum chemistry study. J Phys Chem A 2009, 113 (10), 2092–102.

34. Laage, D.; Elsaesser, T.; Hynes, J. T., Water Dynamics in the Hydration Shells of Biomolecules. Chem Rev 2017, 117 (16), 10694–10725.

35. Scheiner, S., Assessment of the Presence and Strength of H-Bonds by Means of Corrected NMR. Molecules 2016, 21 (11).

36. Scheiner, S.; Gu, Y.; Kar, T., Evaluation of the H-bonding properties of CH?O interactions based upon NMR spectra. Journal of Molecular Structure: THEOCHEM 2000, 500 (1), 441–452.

37. Vallurupalli, P.; Bouvignies, G.; Kay, L. E., Studying “invisible” excited protein states in slow exchange with a major state conformation. J Am Chem Soc 2012, 134 (19), 8148–61.

