## Supplementary Information for "Inverse PI by NMR: Analysis of Ligand ^1^H-Chemical Shifts in the Protein-Bound State"

**Supplementary Material for Inverse PI by NMR: Analysis of Ligand ^1^H-Chemical Shifts in the Protein-Bound State**

**Table 1: Ligand 1 – Experimental vs. theoretical chemical shifts and geometric parameters**

| Proton # | ^1^H-CSP [ppm]  (experimental) | (Δσ) [ppm]  (theoretical) | Compensated? | H-Bond acceptor | Comment |
| --- | --- | --- | --- | --- | --- |
| 1 | -0.33 | 0 | YES | 2.5Å to Gln316 C**O** | CH-O hydrogen bond |
| 2 | -0,94 | 0 | partially | 2.9Å to Val278 C**O** | CH-O hydrogen bond, highly acidic proton; non-optimal H-bond geometry to Val278 |
| 3 | -1,68 | -2,09 | YES | 2.8Å to Tyr281 ring center | CH-π hydrogen bond, good stacking quality |
| 4 | -0,13 | 0 | YES | 2.7Å to Gln367 **O**ε | CH-O hydrogen bond |
| 5 | -0,02 | 0 | - | H_2_**O** | solvent contact maintained  in the bound state |
| 6 | -1,74 | -0,6 | partially | 3.6Å to Phe312 ring center | CH-π hydrogen bond, moderate stacking quality |
| 7 | -0,1 | 0 | - | H_2_**O** | solvent contact maintained  in the bound state |
| 8 | -0,35 | 0 | - | H_2_**O** | solvent contact maintained  in the bound state |
| 9 | -0,1 | 0 | - | H_2_**O** | solvent contact maintained  in the bound state |
| 10 | -1,45 | -0,46 | partially | Trp284^(1^*^)^ | CH_3_-π hydrogen bond, moderate stacking quality |
| 11 | -0,7 | -0,29 | partially | 4.2Å to Phe312 ring center | CH-π hydrogen bond, moderate stacking quality |
| 12 | -0,7 | -0,4 | partially | 4.2Å to Tyr281 ring center | CH-π hydrogen bond, moderate stacking quality |

Table 1: [^1^H-CSP]: NMR derived ^1^H ligand CSP values. [Δσ]: Calculated ring isotropic shielding from X-Ray conformational data using the Pople formalism. [Compensated] refers to the presence of hydrogen bond acceptors for ligand CH and CH_3_ groups respectivly. “YES“ refers to good CH-π or CH-O geometries and “partially” to sub-optimal ones. “H-Bond acceptor” lists the acceptor type and distance extracted from X-ray. (^1^*): NMR CSP for methyl protons 10 was calculated as average of the three individual proton shielding values.

***Calculation of aromatic ring isotropic shielding constant (Δσ)***

In order to calculate the expected shielding exerted by aromatic ring systems, we make use of the Pople model.^1^ Here, the center of the aromatic ring is treated as a point-dipole inducing a magnetic field given by the standard dipole equation:

$\Delta\sigma={10}^{6}\times\frac{{{ne}^{2}a}^{2}}{{4\pi mc}^{2}}\times\frac{{3cos}^{2}\theta-1}{r^{3}}$

Δσ is the change in the isotropic nuclear shielding constant in ppm, n is the number of circulating electrons (n = 6 for all ligands discussed in this work), e is the elementary charge in Franklin, a is the radius of the aromatic ring (1.39 x 10-08 cm), m is the electron mass in gram, c is the speed of light in cm/s. θ is the angle between the ring normal through the aromatic center and the proton to ring center vector in rad. r is the distance from the proton to the ring center in cm.^2^

***[^1^H-^1^H]-NOESY signal assignment***

A scheme of the signal assignment strategy is outlined in Figure 1 of the main text. Resonances of the free ligand can be determined from a simple ^1^H-1D (Figure 1a / red ^1^H-1D) while signals of the bound state are observed when the ligand to protein ratio is adjusted to a 1:1 ratio (Figure 1a / blue ^1^H-1D). The nanomolar affinity of Ligand 1 ensures full ligand saturation. For the analysis of the 2D NOESY spectra a 2:1 ratio of ligand to protein is chosen resulting in the emergence of exchange cross peaks between the free and the bound ligand, making the signal assignment of the bound ligand form via cross peak connectivities a simple task. Signal assignment of the bound state is facilitated by the fact that typically only exchange (EXSY) peaks are observed between the free and bound states (solid black square in Figure 1). As stated before, EXSY cross peaks (solid black) arises from exchange between the free and the bound ligand species while the regular NOESY cross peak (dotted black) originate from spatial proximity. Only in cases where intra-ligand NOEs are already visible in the protein free (apo) form, additional NOESY peaks are observed for the 2:1 mixture. Besides the EXSY cross peak (solid black) an additional exchange-relayed NOESY cross peak connects the free and bound state (dotted black). For Ligand 1 this is the case for neighboring protons (resonances 1&6 and 3&5). Since all NOE cross peaks observed for the NSD3/Ligand 1 system involving free ligand species have a positive sign, the transfer of magnetization must happen via a relayed NOE effect, where the ligand retains an effective correlation time resembling the bound state. Discrimination between the two types of cross peaks is straightforward shall be discussed with the example of proton 1 of Ligand 1 (Figure 1a). The free signal of Proton 1 is connected to three other signals via cross peaks in the NOESY experiment. Strong signal intensities are found for the EXSY signal connecting 1_free_ to 1_bound_ and the NOESY signal connecting 1_free_ and 6_free_. The only other signal connecting a free to a bound signal is the relayed NOESY signal 1_free_ to 6_bound_. The much larger signal intensity of the exchange signal over the relayed NOE signal makes the identification of the bound signal of each individual proton straightforward. Interestingly and important to note, bound ligand resonances that are close to the detection limit in the ^1^H-1D can still be detected via NOESY cross peak signals.

**General Methods**

**Sample Preparation**

Sequence information as well as expression and purification protocols for the PWWP1 domain of NSD3 are given in Böttcher et. al.^3^. In order to obtain deuterated protein, the M9 minimal medium was supplemented with ^15^NH_4_Cl (0.5 g/l) and deuterated glucose (2.5 g/1) dissolved in D2O. After a series of pre-cultures (LB - overnight, 1:1 LB+M9 - 8h, M9 - overnight) containing ampicillin (100 µg/l), M9 medium was inoculated, induced with IPTG (0.25 mM) at an OD600 of 0.4 and grown for 36 h at 20 °C. The final buffer for size-exclusion column contained: 50mM Na-phosphate, 100 mM NaCl, 1 mM TCEP, pH 7.2 – dissolved in D2O.

**Protein NMR spectroscopy**

All protein NMR experiments were conducted at 298 K on a Bruker Avance 600 MHz spectrometer equipped with a TCI cryoprobe. [^1^H-^1^H]-NOESY experiments were acquired using the pulse sequence ‘noesyphpr‘ of the Bruker library. NSD3-PWWP1 sample concentrations were 200 μM. Ligand concentrations were 400 μM for Ligand 1 (2:1) and 1mM for Ligand 2 (5:1). The NOESY mixing time was adjusted to 300 ms. Spectra were recorded using 256 (*t1*) x 2048 (*t2*) complex points with acquisition times of 19.3 ms (*t1*) and 155 ms (*t2*). A total of 32 scans were recorded per t1 increment with a recycle delay of 2 sec.

**References:**

1.	Pople, J., Proton magnetic resonance of hydrocarbons. *The Journal of Chemical Physics* **1956,** *24* (5), 1111-1111.

2.	Platzer, G.; Mayer, M.; Beier, A.; Bruschweiler, S.; Fuchs, J. E.; Engelhardt, H.; Geist, L.; Bader, G.; Schorghuber, J.; Lichtenecker, R.; Wolkerstorfer, B.; Kessler, D.; McConnell, D. B.; Konrat, R., PI by NMR: Probing CH-pi Interactions in Protein-Ligand Complexes by NMR Spectroscopy. *Angew Chem Int Ed Engl* **2020,** *59* (35), 14861-14868.

3.	Bottcher, J.; Dilworth, D.; Reiser, U.; Neumuller, R. A.; Schleicher, M.; Petronczki, M.; Zeeb, M.; Mischerikow, N.; Allali-Hassani, A.; Szewczyk, M. M.; Li, F.; Kennedy, S.; Vedadi, M.; Barsyte-Lovejoy, D.; Brown, P. J.; Huber, K. V. M.; Rogers, C. M.; Wells, C. I.; Fedorov, O.; Rumpel, K.; Zoephel, A.; Mayer, M.; Wunberg, T.; Bose, D.; Zahn, S.; Arnhof, H.; Berger, H.; Reiser, C.; Hormann, A.; Krammer, T.; Corcokovic, M.; Sharps, B.; Winkler, S.; Haring, D.; Cockcroft, X. L.; Fuchs, J. E.; Mullauer, B.; Weiss-Puxbaum, A.; Gerstberger, T.; Boehmelt, G.; Vakoc, C. R.; Arrowsmith, C. H.; Pearson, M.; McConnell, D. B., Fragment-based discovery of a chemical probe for the PWWP1 domain of NSD3. *Nat Chem Biol* **2019,** *15* (8), 822-829.
